# Super-resolution stimulated Raman Scattering microscopy with A-PoD

**DOI:** 10.1101/2022.06.04.494813

**Authors:** Hongje Jang, Yajuan Li, Anthony A. Fung, Pegah Bagheri, Khang Hoang, Dorota Skowronska-Krawczyk, Xiaoping Chen, Jane Y. Wu, Bogdan Bintu, Lingyan Shi

**Affiliations:** Dept. of Bioengineering, University of California San Diego, La Jolla, CA92093; School of Medicine, University of California, Irvine, CA 92697; The Ken & Ruth Davee Dept. of Neurology, Northwestern University, Chicago, IL 60611

**Keywords:** stimulated Raman, super-resolution, deconvolution, multiphoton fluorescence, live tissue imaging, live cell imaging, metabolic imaging, localization microscopy

## Abstract

Unlike traditionally-mapped Raman imaging, stimulated Raman scattering (SRS) imaging achieved the capability of imaging metabolic dynamics and a greatly improved signal-noise-ratio. However, its spatial resolution is still limited by the numerical aperture or scattering cross-section. To achieve super-resolved SRS imaging, we developed a new deconvolution algorithm – Adam optimization-based Pointillism Deconvolution (A-PoD) – for SRS imaging, and demonstrated a spatial resolution of 52 nm on polystyrene beads. By changing the genetic algorithm to A-PoD, the image deconvolution process was shortened by more than 3 orders of magnitude, from a few hours to a few seconds. By applying A-PoD to spatially correlated multi-photon fluorescence (MPF) imaging and deuterium oxide (D_2_O)-probed SRS (DO-SRS) imaging data from diverse samples, we compared nanoscopic distributions of proteins and lipids in cells and subcellular organelles. We successfully differentiated newly synthesized lipids in lipid droplets using A-PoD coupled with DO-SRS. The A-PoD-enhanced DO-SRS imaging method was also applied to reveal the metabolic change in brain samples from Drosophila on different diets. This new approach allows us to quantitatively measure the nanoscopic co-localization of biomolecules and metabolic dynamics in organelles. We expect that the A-PoD algorithm will have a wide range of applications, from nano-scale measurements of biomolecules to processing astronomical images.

---

Raman imaging is a vibrational spectroscopy technique that measures the scattered light corresponding to the vibration of molecules. When incident light alters the polarizability of a molecule, the wavelength of the scattered signal is changed by the resulting vibrational modes. Although Raman scattering imaging reveals structural information of a molecule based on the wavelength change of this scattering signal, the signal of spontaneous Raman scattering is weak, and it is difficult to achieve high speed imaging. In 2008, stimulated Raman scattering (SRS) was demonstrated with greatly amplified signal intensities and has been widely applied to bioimaging ever since (1-3). About 10 years later, deuterium oxide probed stimulated Raman scattering (DO-SRS) imaging platform was reported with the capability of imaging metabolic dynamics and a greatly enhanced signal-to-noise-ratio (3). However, the spatial resolution of SRS imaging still needs improvement. A variety of super-resolution approaches that are capable of detecting single molecule signals have been developed for fluorescence microscopy (4-6). Such methods can achieve a few nanometers or sub-nanometer resolution (7-9). Recently, several super-resolution SRS techniques have been developed (10-19). Nonetheless, it is still challenging to achieve super-resolved Raman imaging without manipulating the samples, and to preserve the temporal resolution without any labeling or additional physical or chemical treatment.

Image deconvolution is a computational strategy that removes distortion (20). Distortion in optical microscopy results in an image blurred by light diffraction, and this blurring is expressed as a point spread function (PSF). A PSF model and deconvolution method allow us to enhance resolution of microscopic images. Several deconvolution methods, such as compressed sensing stochastic optical reconstruction microscopy (CSSTORM) (21), fast localization algorithm based on a continuous-space formulation (FALCON) (22), and sparse image deconvolution and reconstruction (SPIDER) (23), have been developed to achieve super-resolved images by localization of single fluorescence emitters. These methods successfully enhanced the temporal resolution of localization microscopy. However, these methods cannot localize the emitters in general widefield microscopy images and are not capable of detecting single-molecule signals when images are taken with low-sensitivity sensors.

To overcome these limitations, Martinez et al. (24) developed a deconvolution method to fit the measured data by a superposition of virtual point sources (SUPPOSe). This method approximates a super-resolution image by placing a limited number of virtual emitters on the image and optimizing the position of each emitter. The characteristics of this approach are the fixed total intensity as a certain number and quantization of the intensity in each pixel. SUPPOSe sets the total number of virtual emitters, and each emitter has the same unit intensity. The fixed total intensity prevents virtual emitters from deviating away from the optimized position. Because of this characteristic, the residual images can be removed, such as ring artifact (25), as shown in Extended Data Fig. 1a, b. Additionally, due to the fixed unit intensity, intensities at each pixel can only be multiples of the unit intensity. Finally, the two characteristics of SUPPOSe lead to extremely high sparsity of resulting images, which overcomes the limit of aforementioned sparse deconvolution methods. Applying SUPPOSe to our SRS imaging, however, we found major drawbacks in SUPPOSe including slow processing speed and low precision of signal’ s spatial location, which are technically non-trivial.

To significantly enhance the data processing speed and precision of SUPPOSe, we developed a deconvolution method, named A-PoD (Adam-based Pointillism Deconvolution), that uses Adaptive Moment Estimation (Adam) solver instead of a genetic algorithm for optimization process. The gradient descent algorithm, Adam, removes the randomness in the genetic algorithm and enables us to enhance the spatial precision and shorten the data processing time. We applied A-PoD to SRS imaging, and generated a series of super-resolved images of mammalian cells and tissues, as well as Drosophila brain tissues. These images displayed nanoscopic distributions of protein and lipid in biological samples. We further measured the shapes and sizes of individual lipid droplets (LDs) in Drosophila brain samples and examined the effects of high glucose diet on brain lipid metabolism and the size distribution of LDs. Our A-PoD algorithm achieves super-resolution images with higher spatial precision than existing deconvolution methods and at a markedly enhanced speed for image processing.

## RESULTS

### Main concept of A-PoD (Adam optimization-based Pointillism Deconvelution)

We converted SRS images into super-resolution images using a procedure illustrated in Fig. 1a. First, a specific number of virtual emitters proportional to the overall brightness of the image are placed on an image (X), and a blurred image (S) is created through convolution of X and the PSF. When the position of each virtual emitter is adjusted such that the difference between the blurred image S and the measured image (Y) is minimized, X becomes the image with the most optimal distribution of virtual emitters. We used a modified Adam solver (71) for the optimization in A-PoD (see Methods section for details). Using simulation image data, we compared A-PoD with Deconvolutionlab2 with Richardson-Lucy method, a widely used deconvolution algorithm. A-PoD outperformed deconvolutionlab2 using Richardson-Lucy algorithm (even with 100 iterations; see extended Data Fig 1).

**Fig. 1.**
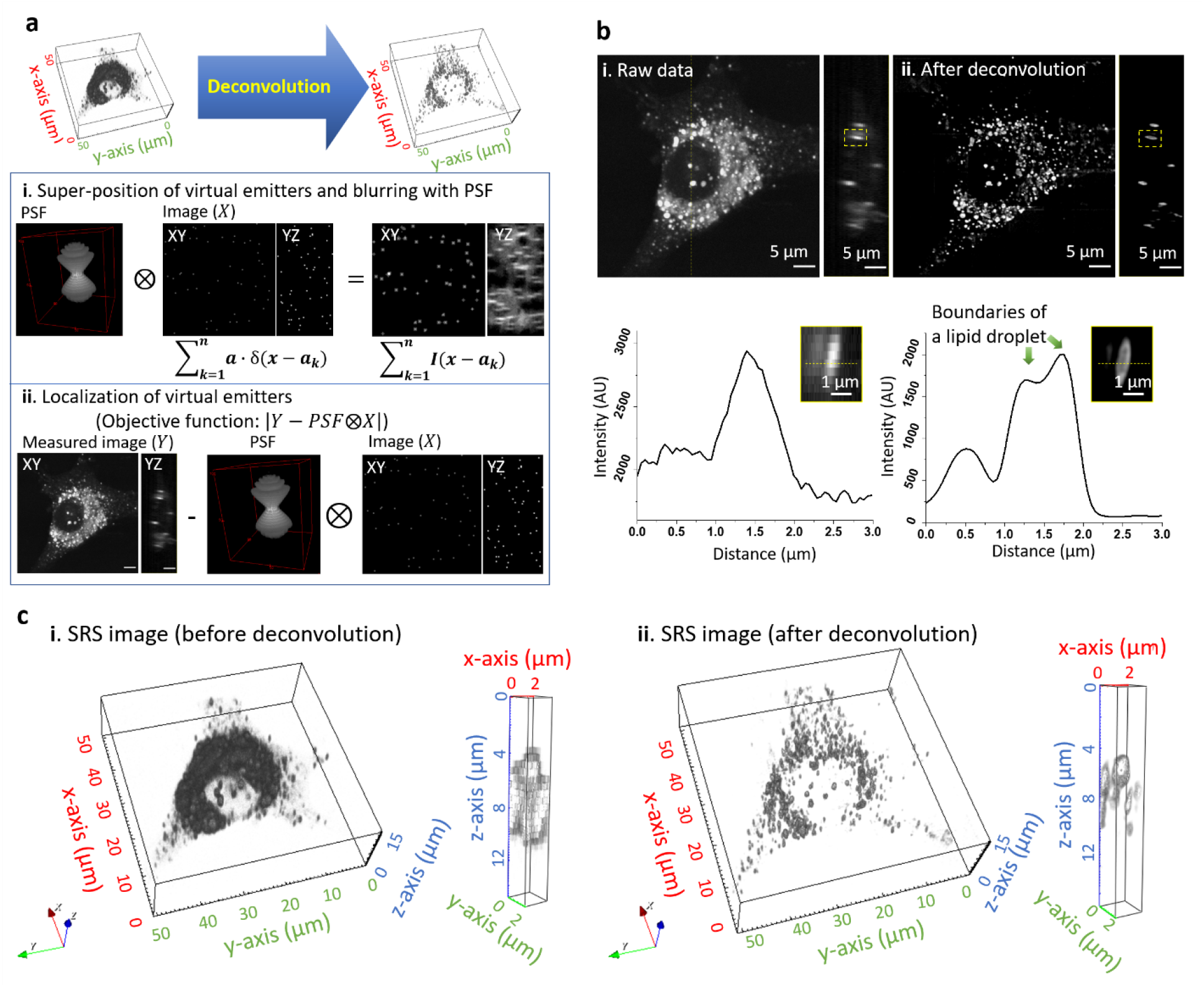
Deconvolution of SRS images using A-PoD. **a**. Schematic of super-resolution SRS image processing. **b**. 3D deconvolution result of lipid droplets (2850 cm^-1^) in a live cell. Following deconvolution, the membrane of an individual LD was clearly visualized in the intensity profile in the lower panel. **c**. 3D rendering results of the SRS image before (i) and after (ii) deconvolution. After deconvolution, the shape of ∼1 μm sized lipid droplets was clearly visible.

As a proof of concept, A-PoD was applied to SRS imaging of lipids (at ∼2850cm^-1^) in a live cell. Application of A-PoD greatly improved spatial resolution (Fig. 1b). The increased spatial resolution clearly revealed individual lipid droplets (LDs) inside the cell, allowing us to distinguish the membrane and the internal space of the LD. The three-dimensional sizes of individual LDs can also be clearly visualized from the sharpened image (Fig. 1c).

To assess the precision of A-PoD, we compared the localization results with those obtained using SPIDER (23). We used a raw mitochondrial image stack in the previous SPIDER publication (23) that was composed of 100 frames. Each image frame contained information of scattered blinking emitters. The image stack was processed with SPIDER program. The widefield image was generated by averaging the stack and was deconvolved using A-PoD. Image processing using A-PoD revealed the mitochondrial structure similar to that obtained using SPIDER (Extended Data Fig 2a, ii and iv). Cross-section signal intensity profiles of images showed that the thickness of the mitochondrial membrane measured by the two methods was almost same. These results demonstrate that A-PoD can reconstruct a super-resolved image from a single frame widefield image. To test the processing speed of A-PoD, we deconvolved the mitochondrial image with limited virtual emitter numbers of 10^5^. As iteration number increased, the similarity between the ground-truth image and the deconvolution image was increased (Extended Data Fig 2b). Using A-PoD, the entire process was completed in 2 seconds, and the similarity was higher than that using genetic algorithm that took 96 minutes with 5×10^6^ iterations. By further increasing the number of iterations, the genetic algorithm could improve the similarity but with a much longer processing time.

### STORM image analysis

To evaluate the spatial precision of deconvolution, we compared A-PoD results with DAOSTORM (26), a widely used algorithm to localize emitters in super-resolution imaging methods such as STORM (Extended Data Fig. 3). For this comparison, an image of cultured neurons where spectrin was labeled using a fluorescent antibody (mouse anti-βII spectrin antibody conjugated with Alexa647) was analyzed using the two algorithms. The original STORM image stack was composed of 16500 frames, and two regions of interest (ROIs) with different emitter densities were analyzed (Extended Data Fig. 3b, c). One of the selected ROIs contained a low emitter density. From the entire image stack of the selected ROI, “epifluorescence”-like image was calculated. The image was deconvolved using A-PoD. Due to the low density of emitters, individual molecules in the image frame could be localized using DAOSTORM. Analysis using either A-PoD or DAOSTORM revealed the periodic structure of the membrane-associated periodic skeleton (MPS) in neurons. The intensity profile and the auto-correlation curves (Extended Data Fig. 3b. iii, iv) showed the periodicity quantitatively, and the periodicity obtained using A-PoD was close to that obtained using DAOSTORM with less than 20% of error. From another ROI containing a high emitter density (shown in the green boxed area in Extended data Fig.3a-i), we analyzed a single frame from the image stack using A-PoD (Extended Data Fig. 3c. I, ii). Interestingly, the periodic structure of the MPS becomes more clearly visible than the other ROI with lower emitter density (Extended Data Fig. 3c. iii, iv). Therefore, A-PoD can be used to analyze images with similar performance as DAOSTORM at low emitter density. A-PoD can be applied to processing images with a wider range of emitter densities.

### Standard sample measurement

To quantitatively determine the resolution of A-PoD coupled SRS imaging, we first analyzed images of standard polystyrene beads with known sizes (270 nm and 1 μm, respectively). The measured image Y was reproduced through convolution of the PSF and the virtual image X. For precise deconvolution, accurate prediction of PSF is critical. We evaluated the results of using PSFs determined by the pump beam (PSF_pump_), the Stokes beam (PSF_Stokes_), and the convolution of PSF_pump_ and PSF_Stokes_ (PSF_conv_ = PSF_pump_ ⊗PSF_Stokes_), respectively. After deconvolution of a 2D image of 270 nm beads and using the decorrelation analysis (25), we obtained the same spatial resolution of 52 nm from all these three approaches, but with different full width half maximum (FWHM) values (Fig. 2a, Extended Data Fig. 4). Using PSF_conv_ reproduced the most accurate bead size, while PSF_pump_ and PSF_Stokes_ showed results approximately 10-20% smaller than the actual size. Therefore, we analyzed the image of 1 μm beads using the PSF_conv_. After deconvolution, the lateral size of the bead was expressed close to 1 μm, but the axial size of the bead was approximately 2.5 times larger. Since the focal volume of a Gaussian beam has a longer shape along the vertical axis (27), the axial resolution is worse than the lateral resolution. Additionally, we observed a cone-shaped afterimage appearing along the optical axis. This is because the direction and intensity of scattering are affected by the size and material of an object, and this scattering behavior is reflected in the shape of the wavefront of light (28, 29). The wavefront of light is distorted by scattering and diffraction. It is difficult to predict using an ideal PSF model. Therefore, the distortion near the bead was not removed by deconvolution. However, this can be mitigated by the combination of adaptive optics and a deep learning method that learns PSF changes around an object (30, 31).

**Fig. 2.**
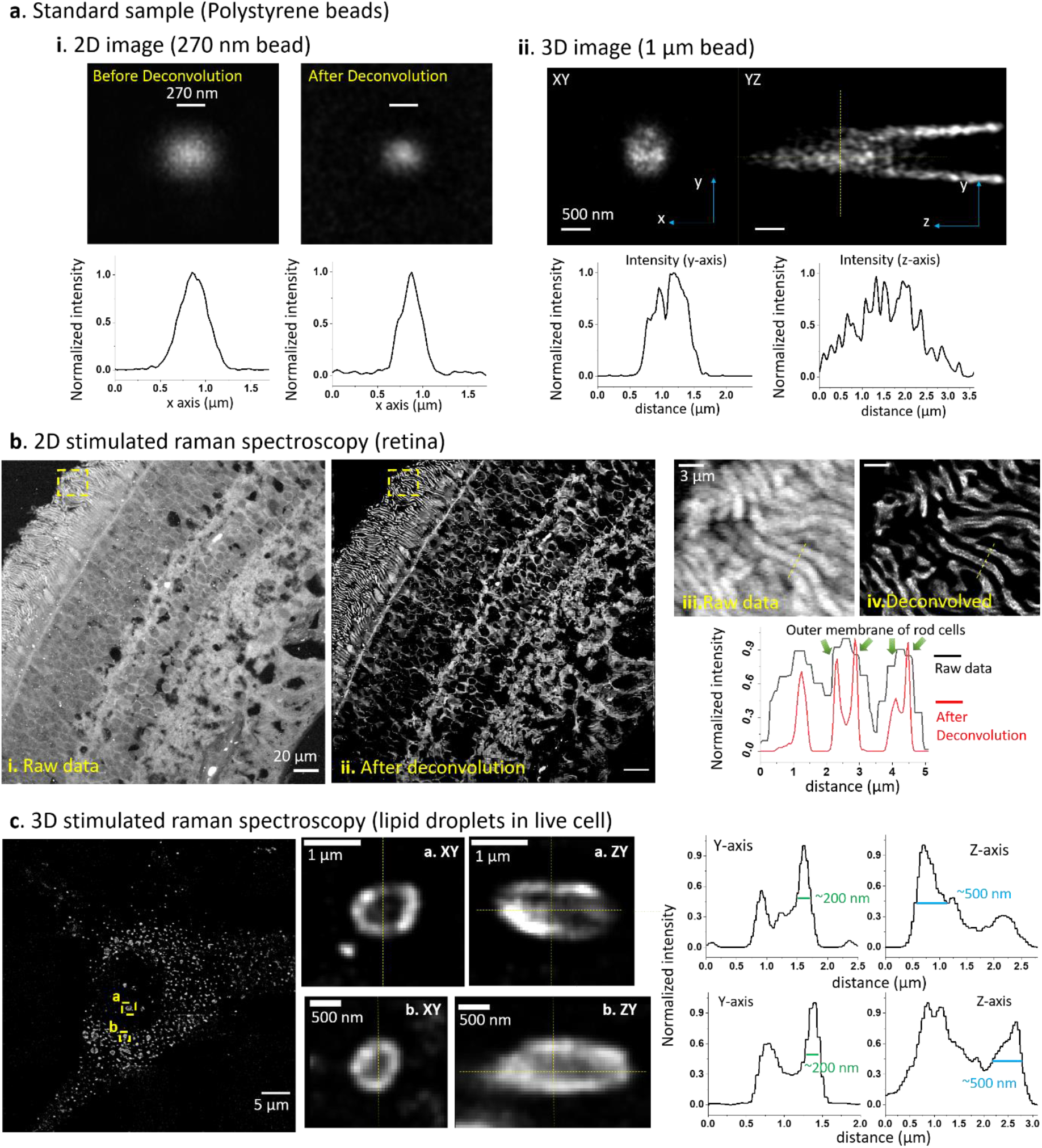
Deconvolution results of SRS images. **a**. Images of standard beads (270 nm and 1 μm beads at 3066 cm^-1^). In panel i, after the deconvolution of a 2D image of a 270 nm bead, FWHM of the intensity profile was decreased from 364 nm to 271 nm. Panel ii shows 3D images of a 1 micron bead before and after deconvolution, together with the corresponding signal intensity profiles. The lateral size of the bead was almost 1 μm, but the axial size was over 2.5 times bigger than lateral size. The tail-like artifact was not removed by A-PoD. **b**. An SRS image of a human retinal section (at 2930 cm^-1^). After deconvolution using A-PoD, contrast of the image was markedly enhanced. Deconvolution results revealed the rod outer segment cell membrane-like intensity profile. The boxed area in the outer segment by the dotted lines in b-i and b-ii is enlarged and shown in b-iii and b-iv.**c**. Deconvolution result of 3D SRS images (2850cm^-1^) lipid droplets in a live cell. Following deconvolution, the detailed structure of LDs was more clearly visualized, including the internal score and the surface membrane.

### Human retinal tissue imaging

Next, we extended A-PoD to SRS imaging of human retinal tissue samples (Fig. 2b). We focused on the outer segments of photoreceptors, which contain membranous photoreceptor discs surrounded by the cell membrane. After applying A-PoD to the SRS image, the image resolution was markedly increased, allowing for improved structural discrimination. For instance, the cell membrane could be visually distinguished in the outer segment of rod cells. The thickness of the cell membrane was about 170 nm, and the resolution of the entire image calculated using the decorrelation analysis method (32) was approximately 100 nm. This resolution with the retinal sample image is lower than the standard bead image, because the deconvolution accuracy depends on the imaging conditions, including the intensity and sampling frequency. The bead image was measured with a sampling rate of 26 nm/px, but the sampling rate of the retinal image was 198 nm/px. Although the increase in spatial resolution was not sufficient to resolve the actual membrane thickness of 4∼5 nm, considering the wavelengths of the laser beams and the characteristics of the PSF, the resolution of ∼100nm clearly exceeded the diffraction limit. It is known that the lipid composition of rod cell membrane is significantly different from the photoreceptor discs (33). The A-PoD coupled SRS microscopy demonstrated a remarkable ability to distinguish these compartments.

### Lipid droplet (LD) imaging

Lipid droplets are organelles important for cell proliferation and survival. These ubiquitous organelles not only serve as energy stores, but also play crucial roles in cell signaling and membrane trafficking. They also contain diverse spatial and chemical information that may reflect oxidative stress, metabolic flux, and disease status (34-44). However, it has been challenging to direct visualize LD metabolism at the organelle level, mainly due to a lack of spatial information in conventional lipidomic modalities. Using A-PoD-coupled DO-SRS imaging, we visualized the nanoscopic distribution of LDs and their metabolic activities. DO-SRS imaging (at 2850 cm^-1^) clearly revealed numerous LDs in the breast cancer cell, and the size of individual LDs could be precisely measured after deconvolution (see Fig. 1c and Fig. 2c). The membrane of each LD was visually separated from the inner space of LD. The thickness of the LD membrane was measured to be approximately 200 nm. This size was similar to the previously measured cell membrane of the rod cells. The axial thickness of the LD was approximately 500 nm, about 2.5 times larger than the lateral resolution. This difference is comparable to the resolution difference in the 1-μm bead imaging.

Next, we used a particle analysis method to remove the background and to focus on the regions of lipid droplets. The subcellular distribution of LDs in the breast cancer cells was then analyzed (Fig. 3). We measured the distances of the detected particles from an arbitrarily chosen point near the center of the nucleus, and calculated the surface area:volume (SA:V) ratio of individual LDs. The LDs were classified into 3 groups based on the distance and SA:V ratio using k-mean algorithm (Fig. 3b). Group 1 had lower SA:V ratio than the other 2 groups (Fig. 3d). The LDs in group 2 (Fig. 3e) were distributed more closely to the nucleus than those in group 3 (Fig. 3f). The capability of A-PoD coupled DO-SRS to identify these different subpopulations of LDs with different SA:V ratio or subcellular distribution may facilitate future studies of dynamic interactions of LDs with other organelles (such as ER), as previous studies suggested that nano-LDs newly detached from ER have high SA:V ratio (76).

**Fig. 3.**
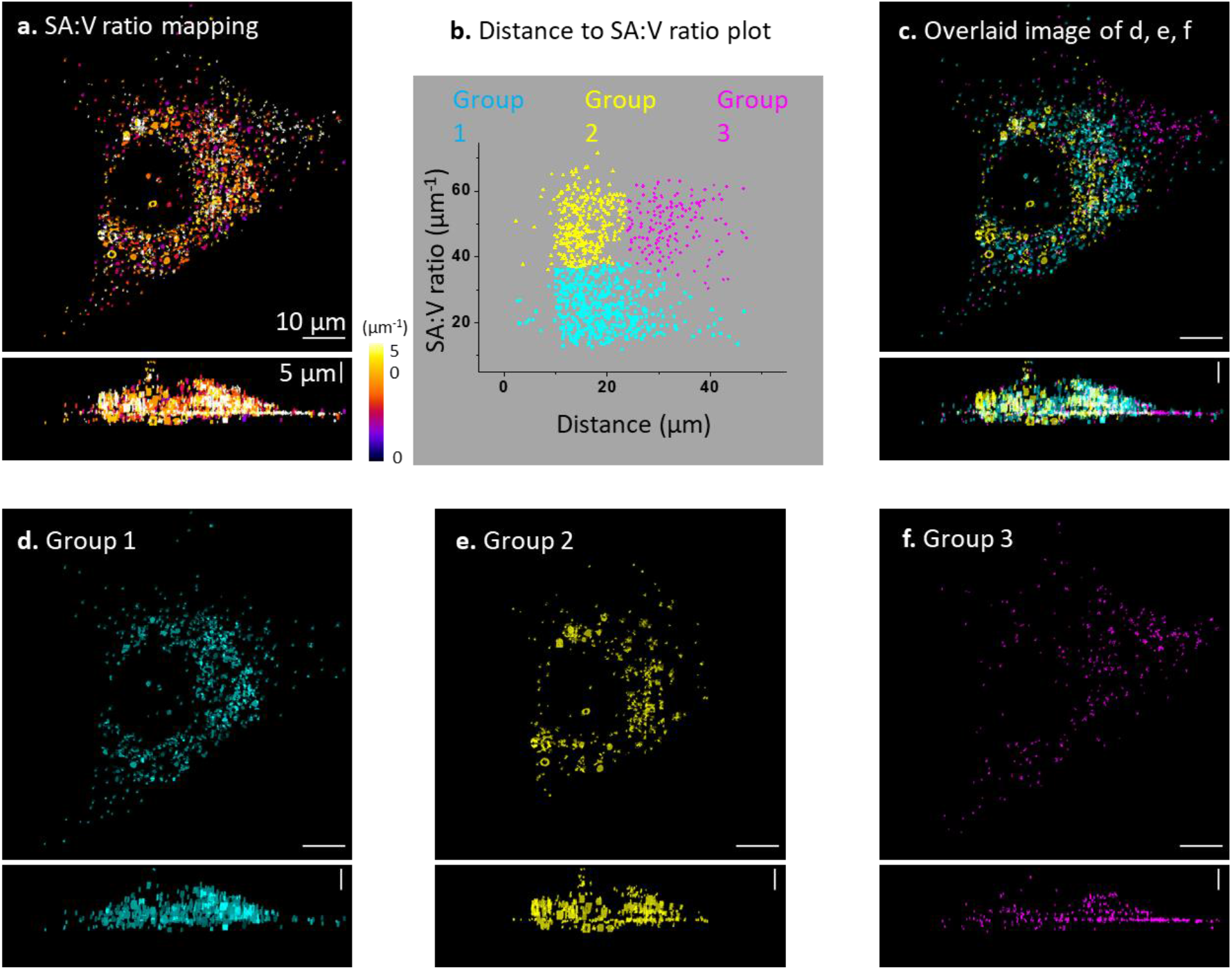
Surface area (SA): Volume (V) ratio analysis. **a**. SA:V ratio of lipid droplets in the breast cancer cell image in Fig. 2c was mapped. **b**. K-mean clustering result shows that the three groups of lipid droplets have different SA:V ratio. **c**. The lipid droplet images in different groups (**d, e, f)** were overlaid. **d**. Lipid droplets in group 1 are widely distributed in the cell, with a low SA:V ratio. **e**. Lipid droplets in group 2 have a high SA:V ratio, and they are distributed closely around the nucleus. **f**. Lipid droplets in group 3 also have a high SA:V ratio, and they were distributed far away from the nucleus.

### Nanoscopic metabolic imaging with super-resolved DO-SRS

Direct visualization of LD metabolism under different conditions at the organelle level is crucial for uncovering the new signaling pathway and molecular mechanisms regulating lipid metabolism. Research in this area has been limited by a lack of spatial resolution in conventional lipidomic imaging modalities. We applied A-PoD coupled DO-SRS metabolic imaging to visualizing lipid metabolism in HeLa cancer cells cultured in the presence of D_2_O. The distribution of LDs in HeLa cells was imaged at 2850 cm^-1^ (CH_2_ vibration) and 2140 cm^-1^ (CD vibration), representing the old lipids and the newly synthesized lipids, respectively (Fig. 4a). After converting the images in each channel to super-resolved ones using A-PoD, the differences in the distribution of old vs. new lipid signals were clearly revealed in 2D (Fig. 4b) and 3D rendered images (Fig. 4e). Thus the metabolic turnover rate of subpopulations of LDs can be quantified with SA:V ratio mapping. Before deconvolution, only areas with concentrated old and new LDs were visualized. After deconvolution, the three-dimensional shape and distribution of individual LDs were clearly visualized. Additionally, we analyzed the surface area and volume of individual LDs from cells cultured under different conditions: the high tryptophan (15x, Trp) and standard control media (Ctrl). The standard deviation of surface area and volume of LDs in HeLa cells cultured in high tryptophan media were wider than those in the control group (Fig. 4c; see Extended Data Fig. 5 for control cells). The SA:V ratio of individual LDs was mapped in deconvolved images (Fig.4d).

**Fig. 4.**
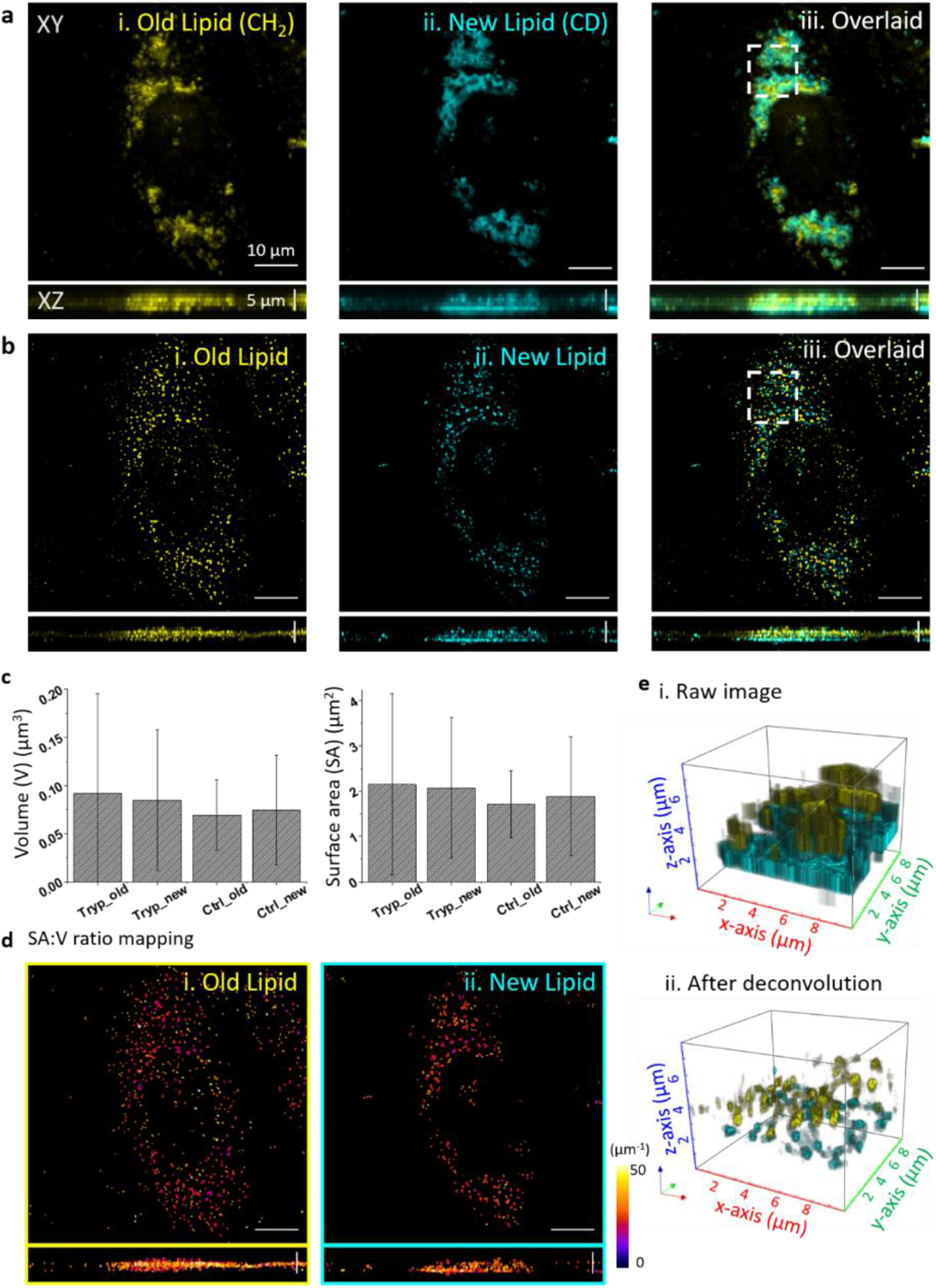
3D super-resolution metabolic imaging of HeLa cell. **a**. DO-SRS images of lipid droplets in CH2 and CD channels. The CH2 channel represents the distribution of old lipid droplets (i), and the CD vibration image shows the distribution of newly synthesized lipid droplets (ii). To compare the two images (i and ii), the images were overlaid (iii). **b**. DO-SRS images were deconvolved using A-PoD, and the results clearly separate the signals of two different types of LDs, old vs. newly-synthesized (i, ii, and iii). **c**. Averaged volume and surface area of each lipid droplets in the two different culture conditions were plotted. The LDs in the cell cultured with excessive tryptophan (Tryp) have wider distribution than those in the control group (Ctrl). The images of the control cell are presented in Extended Data Fig. 4.**d**. The surface area to volume ratio of individual lipid droplets was mapped. Using color code, the SA:V was visualized. **e**. The 3D rendering images of the white dotted boxed regions in panels a (iii) and b (iii) show the resolution difference before and after deconvolution (i and ii).

It has been proposed that LDs play a critical role in the neuroblast cell division and brain development. One major hurdle for understanding functional roles of LDs under physiological or pathological conditions is due to the limited imaging methods for direct observation of LD metabolic activity changes under physiological or pathological conditions. We applied A-PoD-enhanced DO-SRS imaging to directly visualize metabolic changes in Drosophila larval brains collected from animals on different diets. The DO-SRS image of the entire brain lobe collected at 2850 cm^-1^ showed a large amount of lipids in larvae fed with the standard control diet (Fig. 5). To determine the subcellular location of the lipids, the close-up images were taken from the central brain region. These images clearly revealed lipids inside LDs (small dot-like structures). Using A-PoD, we were able to acquire the profile of individual LDs and compare the size distribution of LDs in the brain samples of flies fed standard diet with those fed with high glucose diet (3x glucose) (Fig. 5a-5d; also see histograms in Extended Data Fig. 6a). Size analysis showed that the LDs in 0.2∼0.3 μm^2^ range were predominant in the control group, whereas the LDs in the high glucose group showed a wider range of size distribution, with many small LDs in 0.1∼0.2 μm^2^ range. To better visualize the subcellular distribution of LDs of different sizes in situ, color coded images were generated to show the distribution of small (0.05∼0.2 μm^2^), medium (0.2∼0.3 μm^2^), and large (0.3∼0.45 μm^2^) LDs, respectively (see images in Extended Data Fig. 6a). Considering the small difference in the two histograms and the pixel size (163 nm) in the raw images, it is worth noting that this A-PoD enhanced SRS approach can measure the LD sizes of a wide range, 0.05∼0.45 μm^2^.

**Fig. 5.**
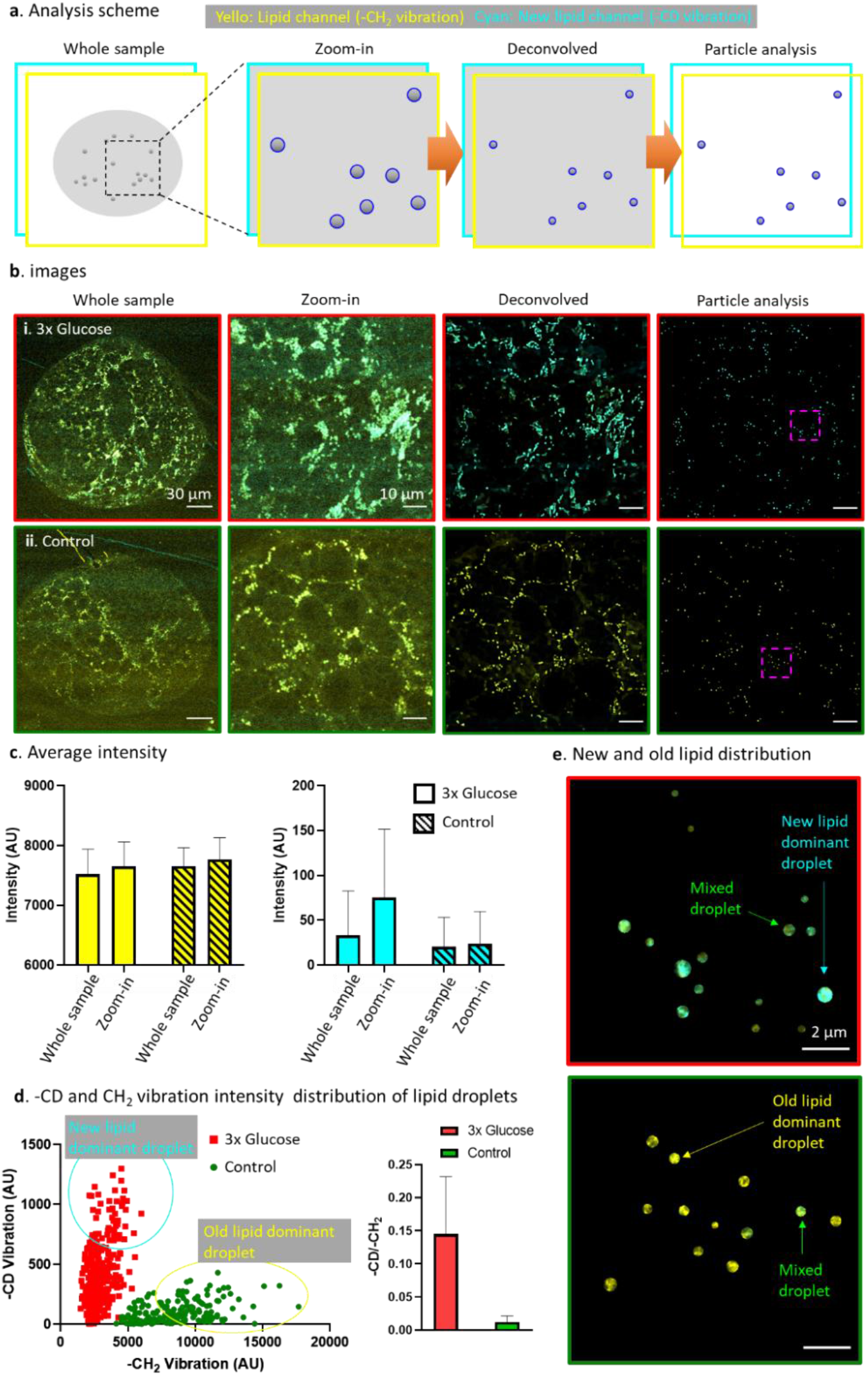
Super-resolution metabolic imaging of Drosophila brain samples. **a**. Schematic of the analysis method. The whole sample image represents the overall lipid distribution. Image was zoomed in to compare the new lipids and old lipids signal distribution of old and new lipids. The nanoscopic distribution of lipids were revealed after deconvolution. The particle analysis method enables us to remove background and analyze individual lipid droplet. **b**. Brain samples from flies on two different diets were measured using DO-SRS microscopy. The sample in 3x glucose group (red boxed images, yellow: -CH_2_ signal, cyan: -CD signal) and control group (green boxed images, yellow: -CH_2_ signal, cyan: -CD signal) were analyzed. The images before the overlay are displayed in Extended Data Fig. 7. **c**. The average signal intensity of the images in two groups. The average signal intensity of old lipid in the control group was slightly higher than the 3x glucose group. The new lipid signal in the 3x glucose group was much higher than the control group. The new lipid signal difference was clearer in zoom-in image. **d**. The scattered plot shows the distribution of new lipid:old lipid (CD/CH_2_) signal ratio of individual lipid droplets. Under the two different dietary conditions, the lipid droplets have clearly distinguishable CD/CH_2_ signal ratio. The averaged turnover rate in the 3x glucose group is over 10 times larger than the rate of the control group. **e**. Using the particle analysis, we can visualize the nanoscopic distribution of newly synthesized lipids in individual LDs. Boxed areas by pink dashed lines in the images in panel b are enlarged and shown.

Combined with D_2_O labeling, the lipid metabolic activities in the brain samples were measured. By measuring the LD size and turnover rates (see Extended Data Fig. 6b), we quantified the correlation between size and metabolic activity. The correlation coefficients were 0.44 in control flies and 0.40 in the high glucose group, with no significant differences detected. Both groups showed a positive correlation between LD size and metabolic activity, suggesting that larger LDs have higher metabolic activity. This result is consistent with our studies on Drosophila fatbody metabolic activity (45-47). Importantly, quantitative analyses of CD/CH_2_ ratio showed that the average lipid turnover rate in the high glucose group was about 10 times higher than that in the control group, suggesting more newly synthesized lipids were accumulated in flies on high glucose diet (Fig. 5d; Extended Data Fig. 6b). The A-PoD coupled DO-SRS combined with particle analysis further enabled us to map distinct subpopulations of LDs: new lipid-dominant, old lipid-dominant and mixed LDs (Fig. 5e). Further studies are necessary to determine molecular mechanisms by which high glucose diet modulates lipid turnover rates.

### Nanoscopic co-localization of proteins, lipids, and fluorophores

Applying A-PoD to spatially correlated multi-photon fluorescence (MPF) imaging and SRS imaging, we next examined nanoscopic spatial distributions of proteins and lipids in mitochondria of live cells (Fig. 6). We imaged HEK293 cells with the mitochondria stably labeled with Mito-Red. The fluorescence signals of Mito-Red were measured using MPF. At the same time, SRS images of 2930 cm^-1^ (CH_3_ protein; in cyan) and 2850 cm^-1^ (CH_2_ lipid; in yellow) were also measured. The SRS images in the two different Raman shifts were unmixed into protein channel and lipid channel using an existing protocol (Fig. 6b).(3, 48) The images of these three measured channels were then converted into super-resolved images using A-PoD (Fig. 6c). Before deconvolution, there was a significant overlap of different types of signals (white areas in Fig. 6d-i). After deconvolution, the white area was reduced, and the different distribution of each component was clearly revealed (Fig. 6d-ii). This is consistent with the fact that SRS signals for protein and lipid panels are not mitochondrion-specific proteins or lipids. On the other hand, the majority of Mito-Red signals (in magenta) were overlapping with lipid signals (in yellow), consistent with the fact that Mito-Red marked the mitochondrial membrane. Furthermore, in the signal intensity profile of the cross-section, the influence of the blurry background signal was reduced after deconvolution, and the position of each component was accurately expressed (Fig. 6d-iii and iv). These data showed that applying A-PoD to multiplexed MPF-SRS imaging significantly enhanced the resolution.

**Fig. 6.**
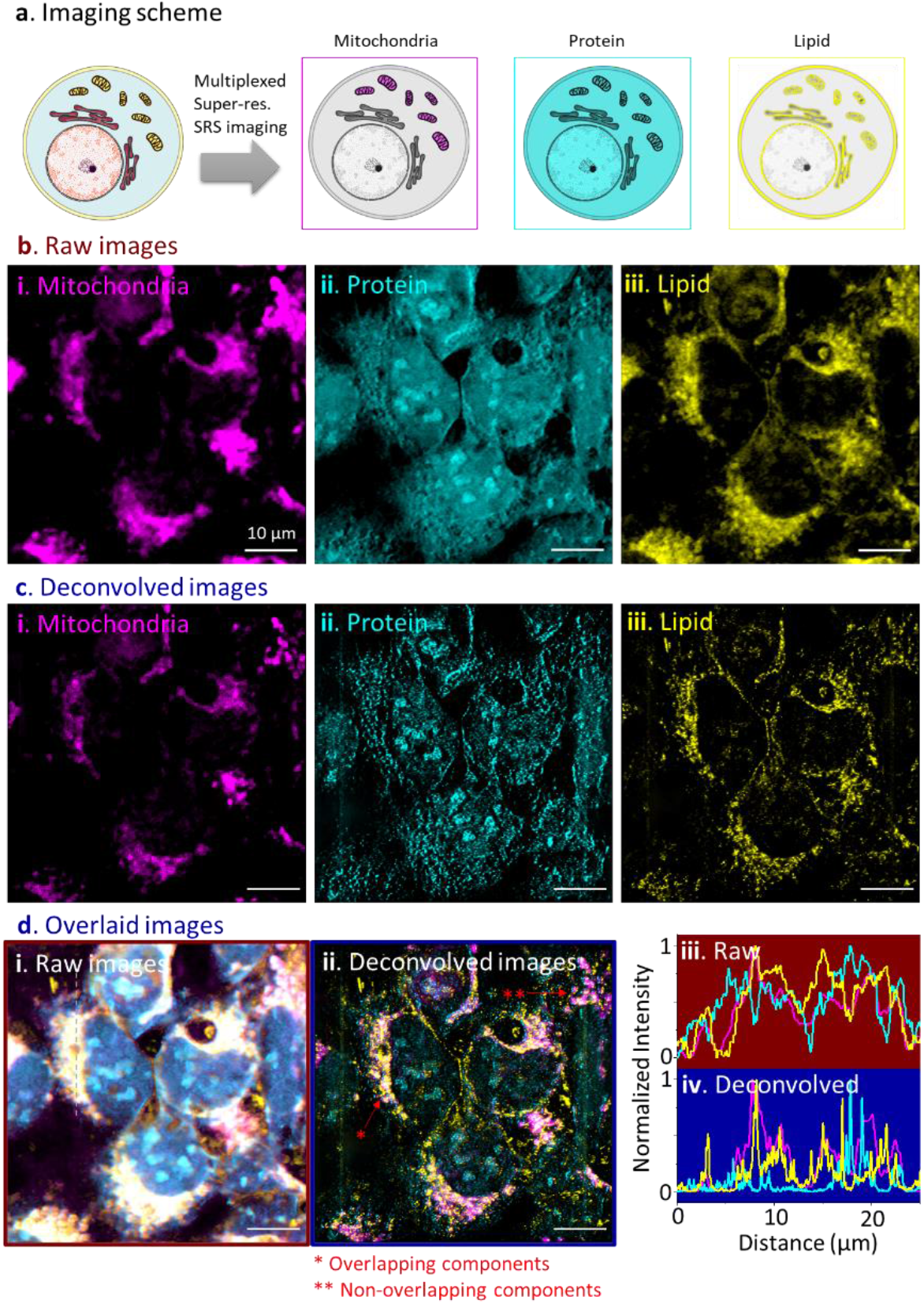
Multiplexed super-resolution DO-SRS imaging of mitochondria. **a**. A The multiplexed imaging scheme. Images were taken in MPF channel (mitochondria) and two SRS channels (protein, lipid) simultaneously, and then deconvolved using A-PoD. **b**. Mitochondria in HEK293 cells were labeled with Mito-Red (magenta) and imaged using MPF and SRS microscopy. The SRS images in 2930 cm^-1^and 2850 cm^-1^ were unmixed to protein (cyan) and lipid (yellow) channels, respectively. **c**. The multiplexed images of mitochondria were deconvolved and converted to super-resolution images. (magenta: Mito-Red; cyan: protein; yellow: lipid) **d**. The superimposed images show the resolution difference before and after deconvolution. The superimposed image before deconvolution (i). After deconvolution, the white area where the three components were overlapping was much reduced (ii). Panels iii and iv show the normalized signal intensity profiles before and after deconvolution. The three components show distinct spatial distribution after deconvolution as shown in the signal intensity profiles.

## DISCUSSION

In this study, we have developed the A-PoD algorithm and integrated it with SRS, DO-SRS, and MPF-SRS imaging methods. A-PoD significantly enhances the spatial resolution of images at a high processing speed and spatial accuracy when an appropriate PSF is defined, regardless of the imaging modalities. A-PoD can be applied not only to widefield fluorescence microscopy (24) but also to various other microscopy techniques. The super-resolution A-PoD-coupled SRS microscopy introduced here also has broad applications including deep-tissue imaging, hyperspectral imaging, and multiplex imaging (49-52).

We first characterized A-PoD as a sparse deconvolution method by analyzing the simulated data. The capability of A-PoD to generate super-resolved image was evaluated by comparison with localization microscopy data (see Extended Data Fig. 2a). Although the genetic algorithm in SUPPOSe is good to optimize variables in an integer domain (e.g., the address of specific pixels), it contains randomness in the process. Gradient values of the function need to be calculated in every optimization step, which is time consuming. Of note, by changing the genetic algorithm to A-PoD, the image deconvolution process was shortened by more than 3 orders of magnitude, from a few hours to a few seconds (See Extended Data Fig. 2b). Compared with the Richardson-Lucy algorithm (See Extended Data Fig. 1c and 8), the most widely used deconvolution method, A-PoD offers much richer chemical information at a high resolution.

For analysis of STORM imaging data (Extended Data Fig. 3), A-PoD demonstrated the potential as an image processing tool for localization microscopy. Generally, in order to achieve a super-resolved image using STORM, we need to keep a low concentration of emitters. In contrary, for A-PoD, strong signals are desirable to achieve a higher resolution. Due to this unique characteristic of A-PoD, we could clearly visualize the periodic structure of the membrane-associated periodic skeleton in neurons from a single image in the bright ROI. This finding implies that A-PoD significantly improves the temporal resolution of localization microscopy, allowing us to extract image features from single to a few frames rather than analyzing tens of thousands of image frames. Depending on the imaging rate of the image stack, it would be possible to take a super-resolved image in a few micro-second ranges when enough emitter density is secured.

Using A-PoD-coupled SRS microscopy, we successfully examined the distributions of proteins and lipids in cultured cells and tissue samples at the nanoscopic level. The nanoscopic distribution of LDs in cancer cells and the membranous outer segments of rod cells in the retinal tissue were clearly resolved. Furthermore, integration of A-PoD into our DO-SRS platform enabled us to examine different distributions of newly synthesized lipids versus the pre-existing lipids in live cells and tissues. This combination provides a powerful tool for direct visualization of lipid metabolic changes not only in cells but also in brain tissues (Fig. 4 and 5).

Using the cultured HeLa cells and the breast cancer cells, we demonstrated the power of A-PoD-coupled SRS imaging in examining subcellular organelles, such as LDs and mitochondria (Figs. 3, 4, and 6). We mapped the SA:V ratio of individual LDs. Since the accuracy of the measured surface area and volume depends on the spatial resolution of images, A-PoD is a valuable tool for analyzing the exact values of these parameters. Using A-PoD-based DO-SRS, we examined the spatial distribution of distinct subpopulations of LDs, those predominantly containing newly synthesized lipids, those mostly containing old lipids and LDs containing mixed lipids. Mapping the old and new lipid domains in individual LDs (Fig. 5e) provides useful information in studying lipid metabolism at the nanoscale. Future experiments are necessary for understanding the pathophysiological roles of LD heterogeneity. Nevertheless, our A-PoD-based DO-SRS imaging system provides a robust method for studying molecular heterogeneity in living organisms.

Analyses of the LD size distribution and lipid turnover rate in Drosophila brain samples indicate that the subpopulation of LDs with higher turnover rate increased in the brain in flies on a high glucose diet and that average lipid turnover rate in the high glucose group was much higher than the control group. It suggests that smaller LDs, which are usually referred to the newly born LDs connected endoplasmic reticulum (ER) (53), may have lower *de novo* lipid synthesis ability. They may obtain lipid content directly from ER lumen. This is consistent with a previous study (54) reporting that enzymes (such as DGAT2, CCT1)-mediated de novo lipid synthesis were mainly localized in the larger mature LDs detached from ER. Here, our A-PoD enhanced super-resolution DO-SRS imaging has revealed the metabolic diversity of LDs, which had never been reported by using other methods. Previous studies reported that the ER stress was induced by high glucose (55), and that ER stress increased the LD number (56, 57). Our A-PoD-based DO-SRS imaging system provides an effective tool for future studies on dynamics changes in LDs, functional roles of LDs and underlying mechanisms under various physiological and pathological conditions.

To define nanoscopic distribution of different molecules, we can utilize A-PoD in multiplex SRS imaging. We prepared HEK293 cells stably expressing Mito-Red and examined subcellular distribution of mitochondria, proteins and lipids using A-PoD based MPF and SRS imaging. As expected, the majority of Mito-Red signals overlap with CH_2_ lipid (membrane) signals (Fig. 6d).

By comparing the spatial localization of different components, we can clearly define co-localized and non-overlapping components. Furthermore, A-PoD coupled multiplex SRS can be applied to imaging other biomolecules such as nucleic acids, etc..

Recently, various super-resolution techniques have been applied to SRS imaging (11,12,61,62). Nonetheless, the localization method for super-resolution fluorescence microscopy was considered not applicable to Raman imaging. Due to the lack of single-molecule detection capability and the high emitter density, it is challenging to localize every single molecule in Raman image. However, A-PoD allows us to overcome this limit by the localization process of virtual molecules. The potential application of A-PoD in new localization microscopy methods was demonstrated in the analyses of STORM images. For the existing localization microscopy methods, the amount of emitter signals has to be precisely adjusted. To make this adjustment, one needs to take numerous different frames to reconstruct a single super-resolved image. However, A-PoD can maximize the temporal resolution by overcoming the limitation of emitter density. Therefore, this program allows us to take not only a super-resolution SRS image but also a super-resolution fluorescence image at a high speed.

A-PoD has a wide range of applications. In this study, we presented the results combining A-PoD with STORM or DO-SRS or multiplex MPF-SRS. It is also applicable to other imaging techniques, in which blurring kernel can be defined. For example, in the case of atomic force microscopy, the PSF model of optical microscopy cannot be applied because the morphology of the sample is measured by force between the tip end and the sample. However, the tip convolution effect blurs images due to the shape of the tip end. An attempt to deconvolution assuming the shape of the tip end was made two decades ago (58), but it has not yet improved the quality of the AFM image dramatically where A-PoD can be applied as a solution for enhancement. In addition, the resolution of super-resolved images can be further improved by using A-PoD. Structured illumination microscopy (SIM), one of the super-resolution imaging techniques, is an example. Studies on the super-resolved Raman imaging technique using structured illumination microscopy (SIM) were published recently (59, 60). Although SIM improves spatial resolution over two folds by reducing the size of the PSF, the resolution of SIM images can be further increased using A-PoD, because deconvolution is also possible based on the reduced PSF. This approach has been applied using a different deconvolution program, Sparse-SIM (61). Finally, A-PoD can be applied to astronomy (62), which is a research field where deconvolution is widely used. In fact, the Richardson-Lucy algorithm was originally published for astronomy studies (63). Taken together, the results shown in this study represent a beginning of many different applications of A-PoD from the nano-scale to the astronomic scale.

**Extended Data Fig. 1.**
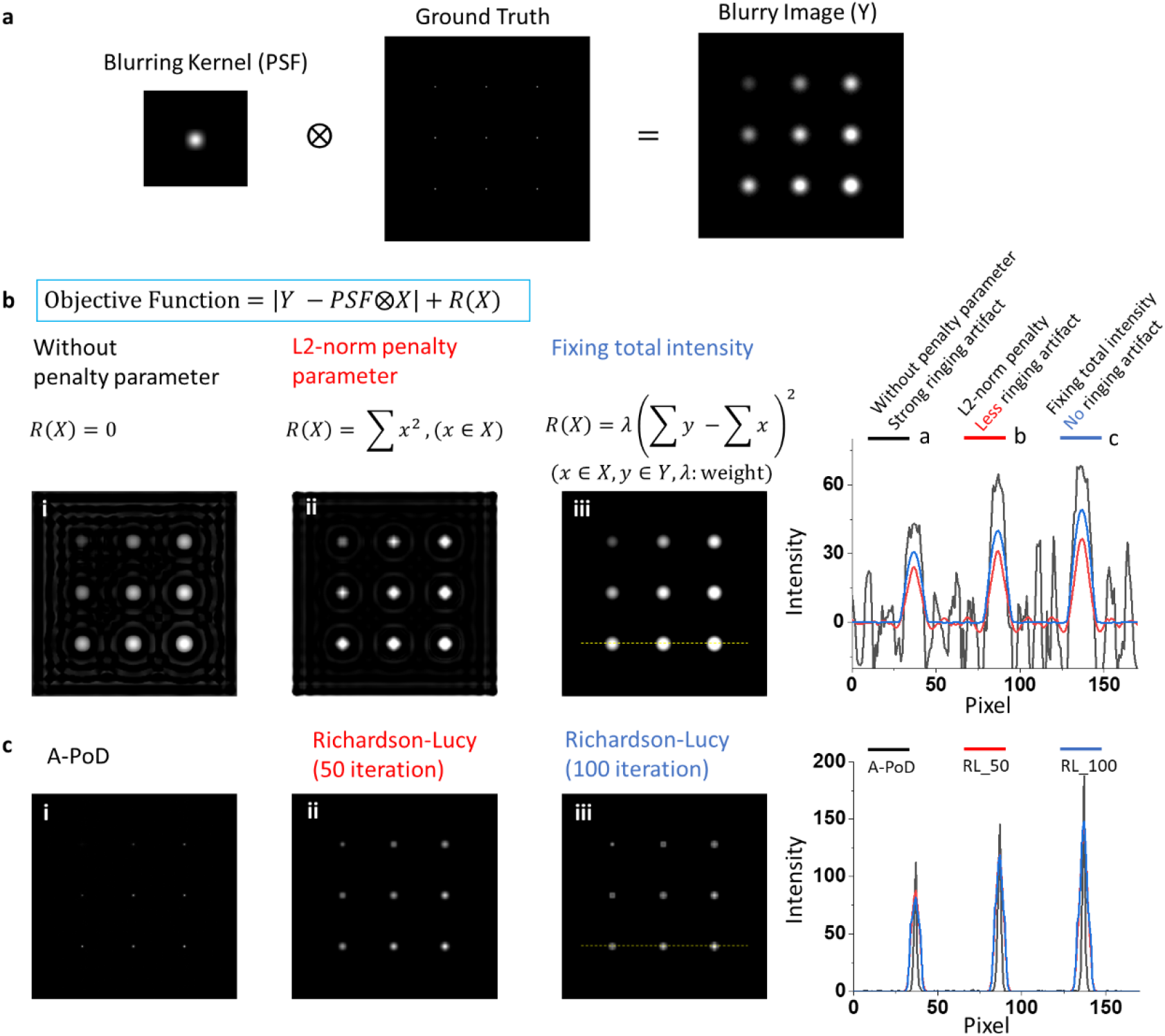
Comparison of A-PoD with Richardson-Lucy method using simulation data. **a**. To compare different deconvolution methods, we generated an artificial image composed of single pixel sized 9 dots. The dots in the image have different intensity values. By convolution with an artificial PSF, a blurry image (Y) was generated. The image (Y) was deconvolved using a penalized regression method. **b**. When we minimize the objective function in panel b, the images, X results. Depending on the penalty parameter, R(X), X has various forms. The optimization result without any penalty parameter has strong ringing artifact as shown in panel b(i), and the result with L2-norm penalty parameter has reduced ringing artifact as shown in panel b(ii). By limiting summation of total intensity, we can reduce the ringing artifact as shown in panel b(iii). The penalty parameter limiting the total intensity as a fixed value makes the values in empty space to zeros. Accordingly, one of the main characteristics of A-PoD, the fixed total intensity of X, can increase sparsity of resulting images. **c**. Comparison of A-PoD with Richardson-lucy method. When we apply another characteristic of A-PoD, quantization of intensity value, together, the resulting image of A-PoD has higher resolution than that obtained using Richardson-Lucy method. The signal intensity profile shows the difference in resolutions. The dots in the A-PoD image have narrower width than Richardon-Lucy images. The calculation time of A-PoD was 1.9 s, and Deconvolutionlab2 using Richardson-Lucy algorithm calculated the image for 1.1 s (50 iteration) and 2.2 s (100 iteration).

**Extended Data Fig. 2.**
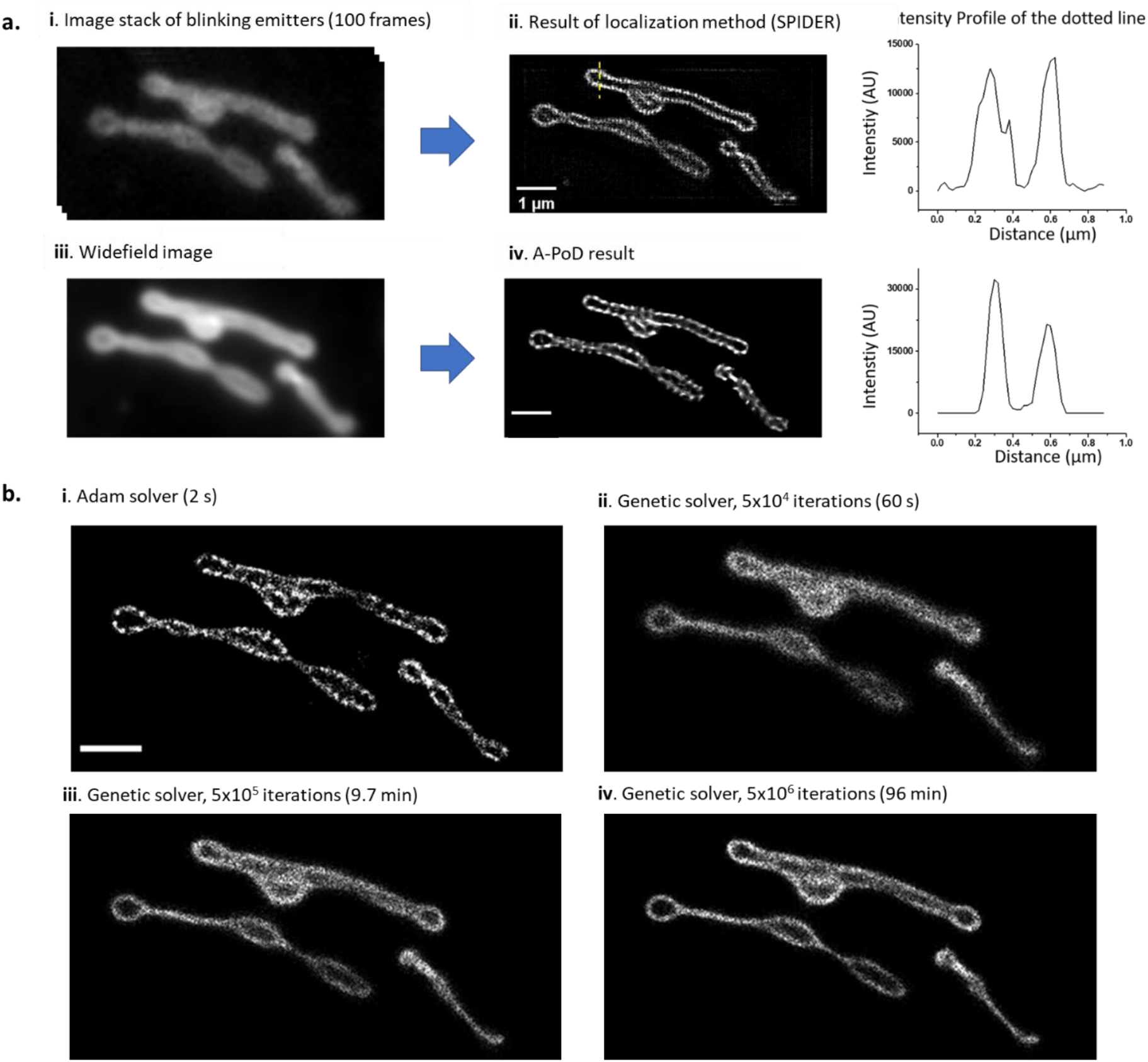
Precision and speed of A-PoD in comparison with SPIDER. **a**. To compare the localization microscopy image with A-PoD result, we deconvolved a mitochondrial image. The image stack is composed of 100 frames. Each image frame contains information about blinking emitters. The emitters were localized using SPIDER deconvolution algorithm. By averaging the image stack, we generated a widefield image, and the widefield image was deconvolved using A-PoD. The intensity profiles of the cross-section in the deconvolved images show the similarity between the two results. **b**. Two optimization methods for the deconvolution process were compared. An image composed of 100000 virtual emitters was deconvolved using the two different optimizers. The results of Adam solver (i) finished calculation within 2 s. By increasing the iteration number, the deconvolution results using genetic solver (ii, iii, and iv with different iteration numbers) were compared with the result of Adam solver. The deconvolution result with a high iteration number shows more precise image. However, to generate an image having same quality as that obtained with the Adam solver, we need to increase the iteration number further beyond 5× 10^6^ more.

**Extended Data Fig. 3.**
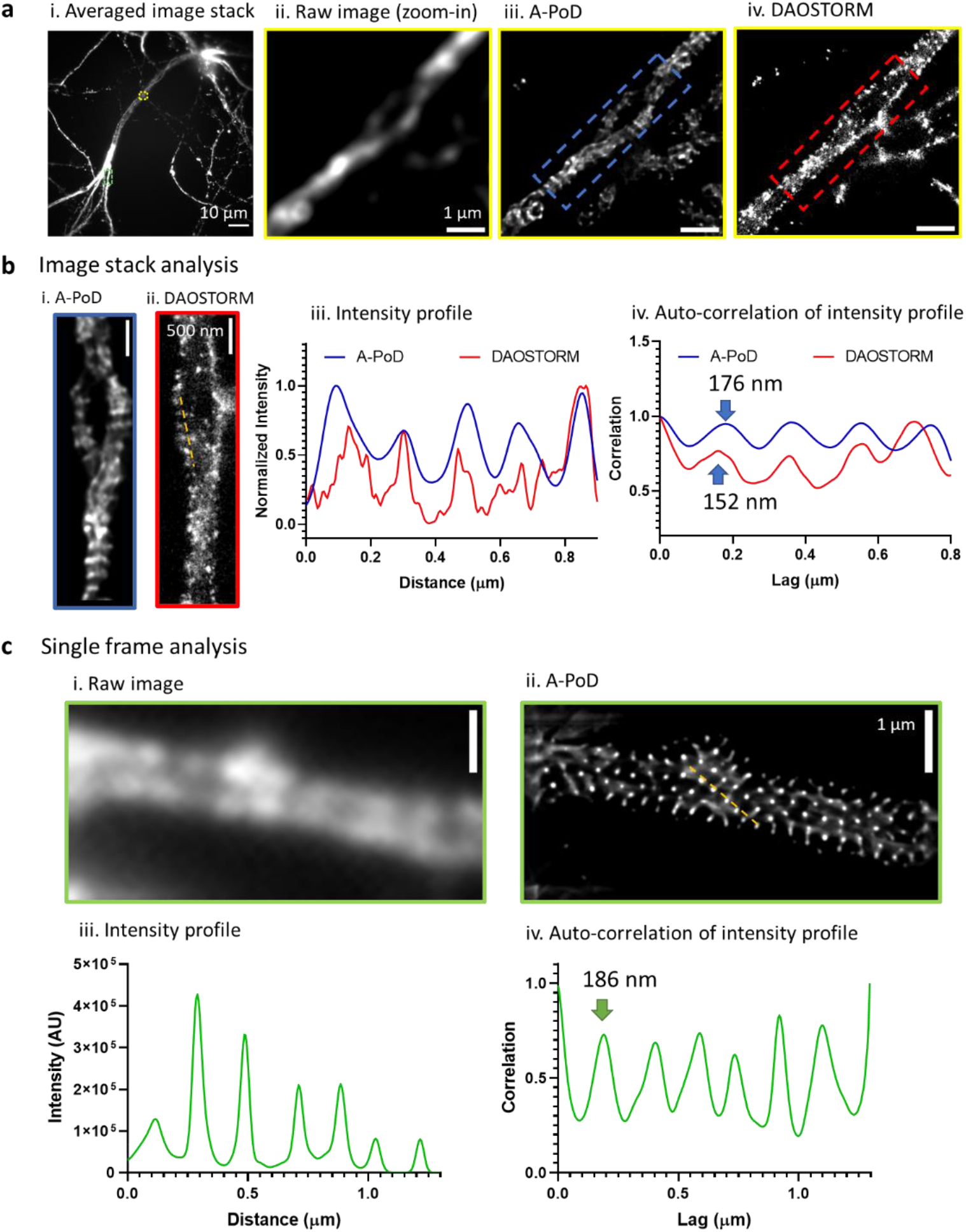
Comparison of the deconvolution results on STORM images using DAO STORM*(26)* versus A-PoD. **a**. (i) A single “epifluorescence”-like image was calculated by averaging the STORM-stack. (ii) We selected an area with low emitter density (yellow rectangle region in (i)) than other areas. (iii) The averaged image stack of the chosen area was deconvolved using A-PoD. (iv) From the whole stack of the selected area, the individual single emitters were localized using DAOSTORM. **b**. The two areas marked by the blue and red rectangle areas in (a. i and b. ii) were selected. (iii and iv) The intensity profiles and auto-correlation data shows the periodicity of the structure of the membrane-associated periodic skeleton (MPS) in neurons. **c**. Another bright area with high emitter density (green rectangle area in a.i) where we cannot localize the individual molecules using DAOSTORM was selected. (i) From the image stack of the selected area, we chose a single frame. (ii) Using A-PoD, we deconvolved the chosen frame. (iii and iv) The intensity profile and the auto-correlation result show the periodicity. Due to the strong intensity, the periodic structure was clearly revealed, and the interval in the MPS is also close to the previous published result, 190 nm (64).

**Extended Data Fig. 4.**
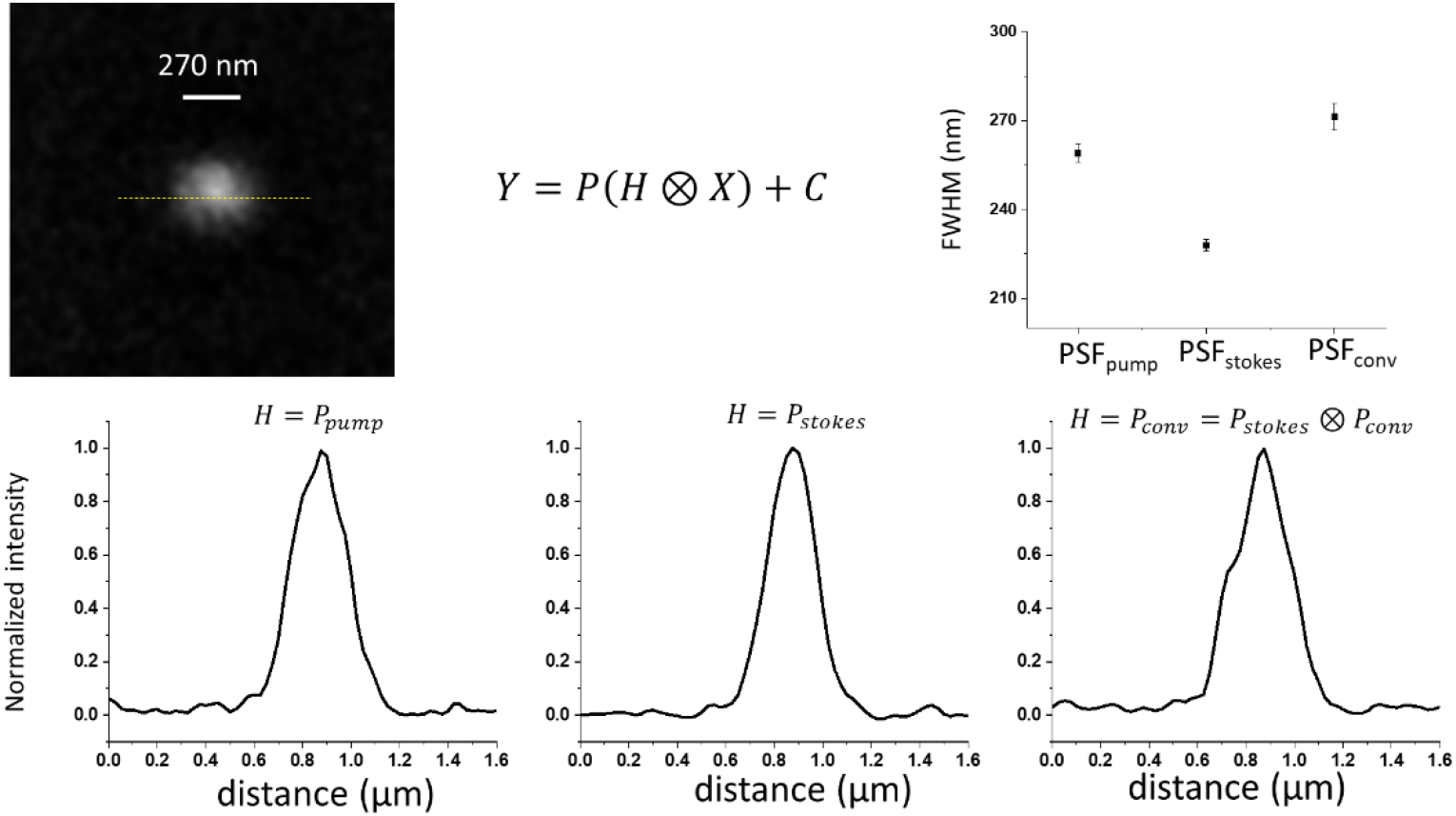
Comparison of three PSF models. The bead image of 270 nm size was deconvolved using different PSF models. To compare the accuracy of the results, we compared the FWHM of the fitted Gaussian functions. When we used PSF_conv_, FWHM was almost same with the bead size.

**Extended Data Fig. 5.**
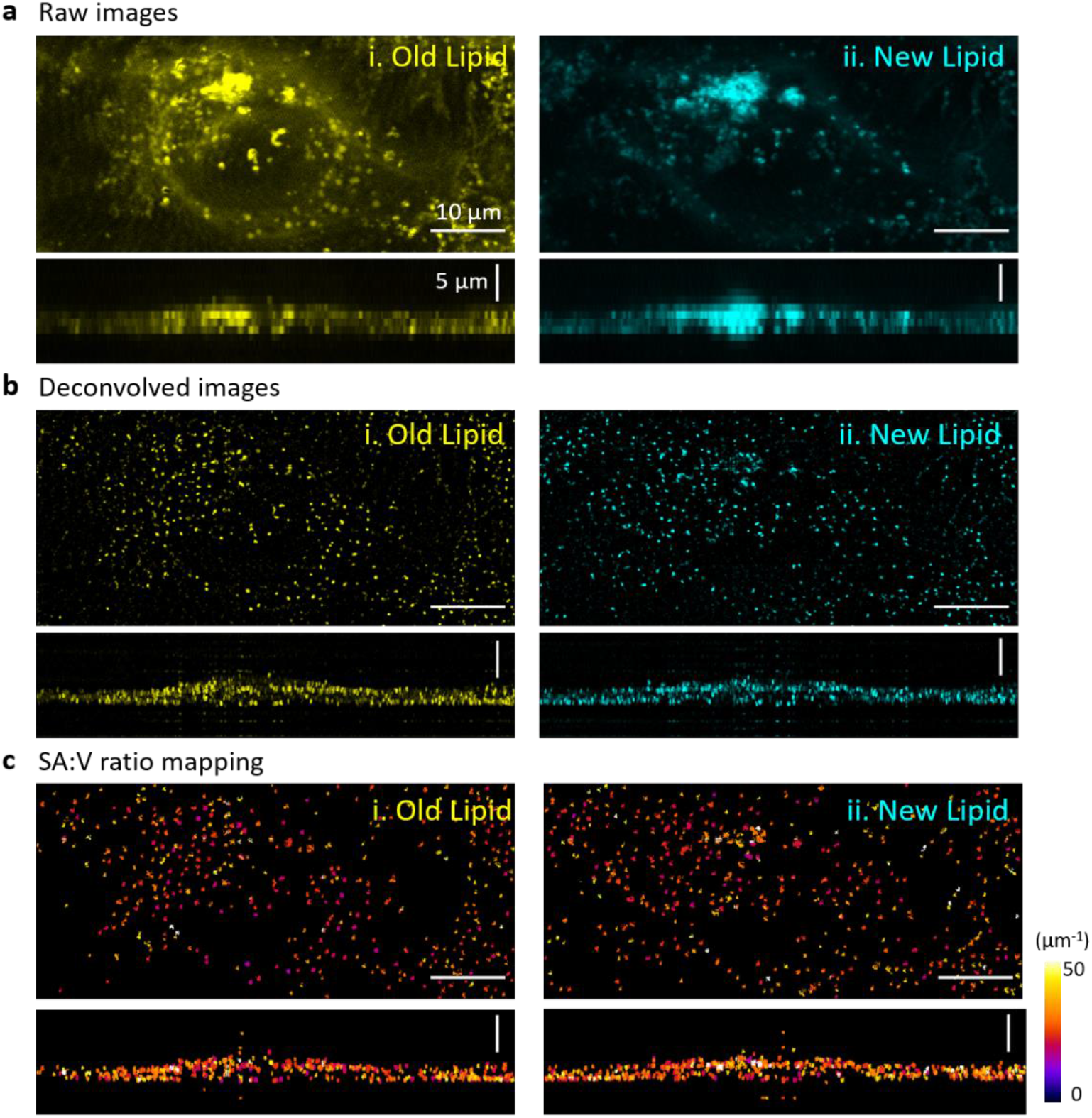
SRS images of a HeLa cell cultured in the standard medium. **a**. Raw DO-SRS images of the HeLa cell. **b**. Deconvolution results of the images. The images show the shape and distribution of the lipid droplets in sub-micron scale. **c**. After measuring the surface area and volume of individual lipid droplets, the surface area to volume ratio of individual LDs was mapped.

**Extended Data Fig. 6.**
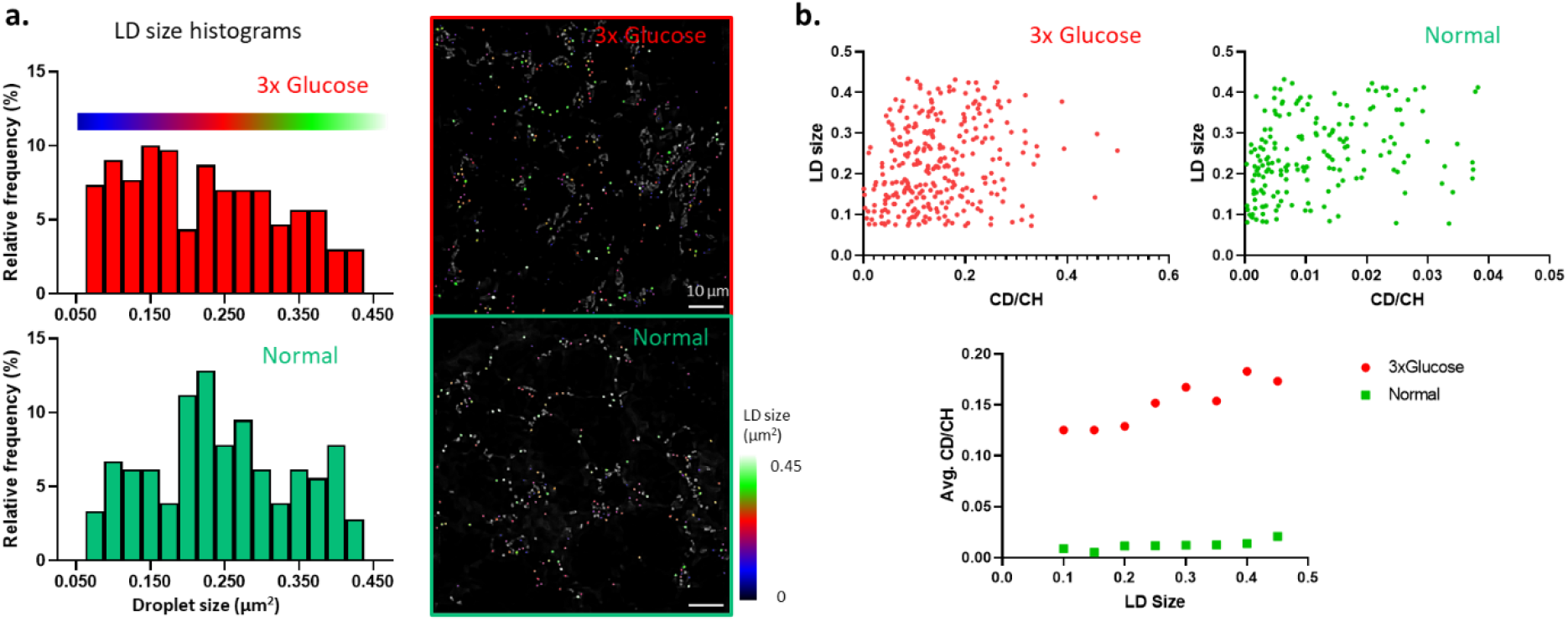
LD size and lipid turnover rate distribution. **a**. In flies fed on different diets, LDs have different size distribution. In high glucose group, the LD size was widely distributed, and the number of LDs in 0.1∼0.2 μm^2^ range was higher than the other size. In control dietary condition, the control group with standard diet, the number of LDs in 0.2∼0.3 μm^2^ range was high. LDs were labeled on the images with three colors according to the size (Blue, 0.05∼0.2 μm^2^; Red, 0.2∼0.3 μm^2^; Green, 0.3∼0.45 μm^2^). **b**. To compare the LD size and lipid turnover rate, the two parameters of individual LDs were plotted. Under both conditions, LD size and lipid turnover rate show positive correlation. Correlation coefficient: 0.40 (3x glucose), 0.44 (control).

**Extended Data Fig. 7.**
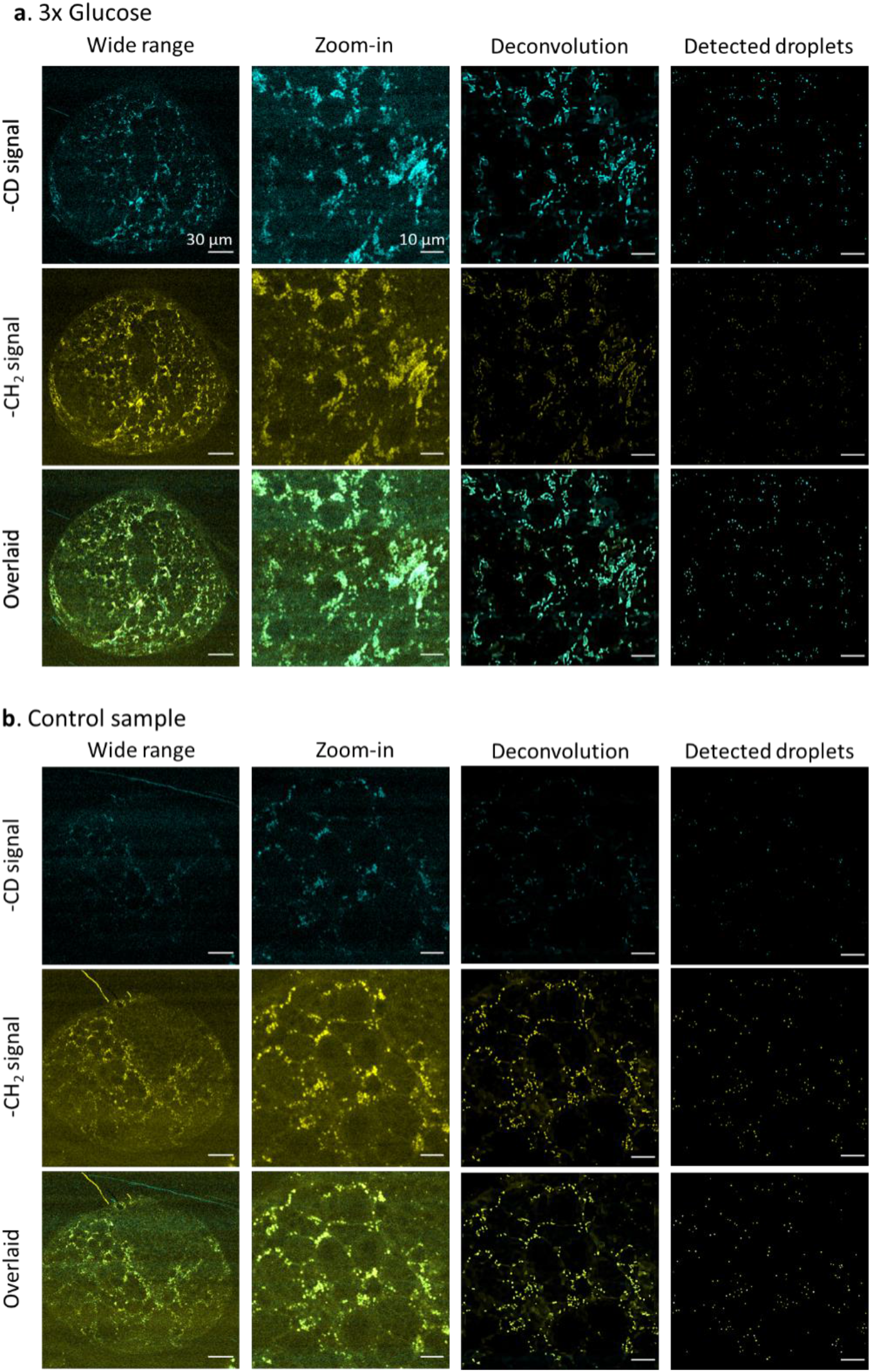
SRS images of larvae brain samples from flies fed on different diets. **a**. DO-SRS images of a drosophila larvae brain in 3x glucose group. The wide range new lipid (CD) and old lipid (CH_2_) signal show the distribution of newly synthesized lipids and old lipids in whole sample, respectively. In the zoomed-in images, the microscopic distribution of two different lipid components is clearly shown. After deconvolution, the nanoscopic distribution and shape of lipid droplets are getting clearer. By using the particle analysis method, we can remove the background and focus on the areas of lipid droplets. **b**. SRS images of a drosophila larvae brain in the control group were processed with the same manner in a. These images were analyzed, and the analysis result is explained in Fig. 6.

**Extended Data Fig. 8.**
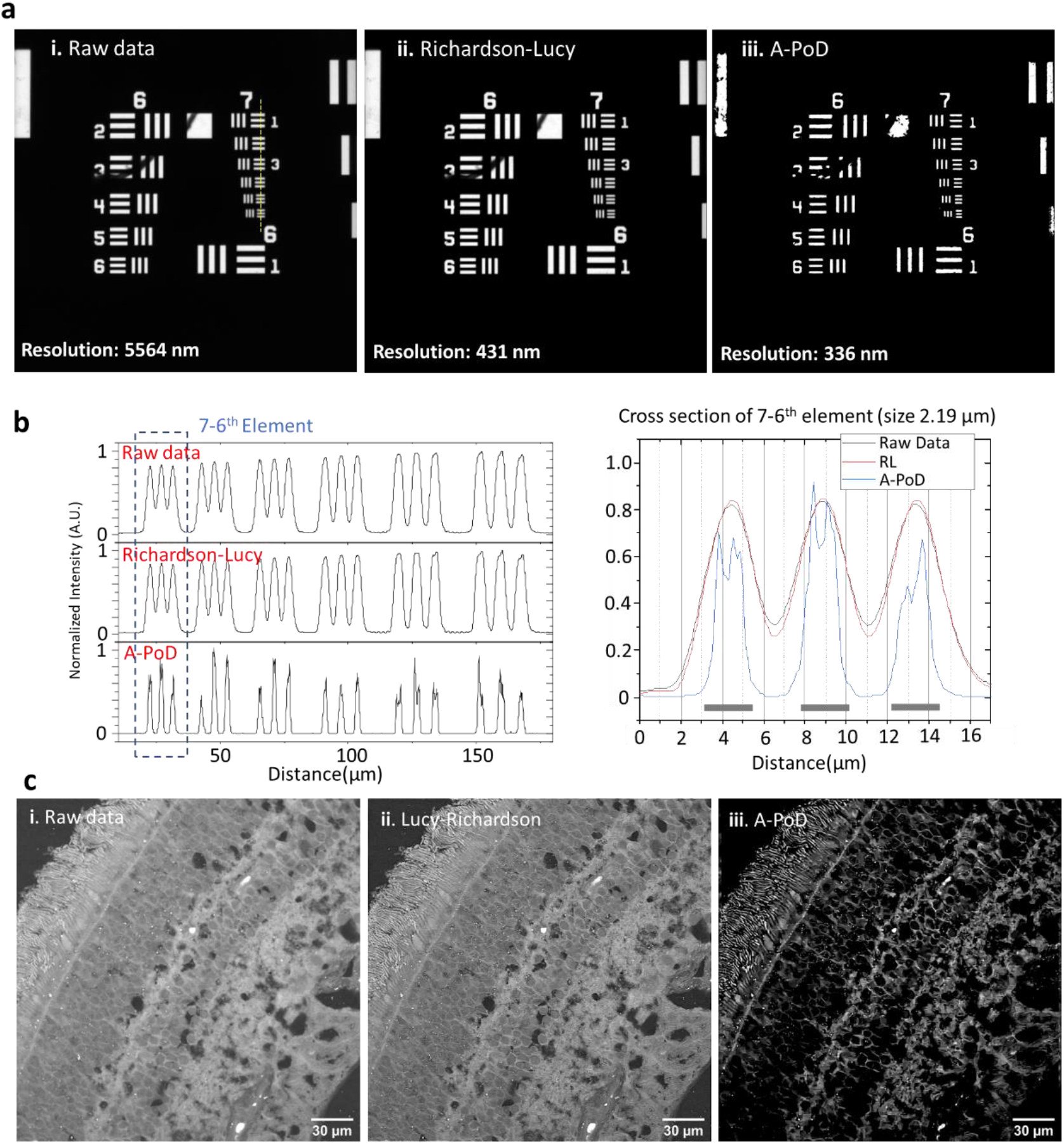
Comparison between A-PoD and the Richardson-Lucy method. **a**. USAF-1951 resolution target. The fluorescence image of the resolution target in the paper(65) was deconvolved using Richardson-Lucy algorithm (Deconvolutionlab2 program)(20). **b**. Intensity profiles of the yellow dotted line in figure A show the resolution difference. A-PoD result resolved each line perfectly, but Richardson-Lucy result could not resolve them. **c**. Deconvolution results of retinal tissue image. The raw image (i) was deconvolved with Richardson-Lucy algorithm (ii) and A-PoD (iii). The image contrast was significantly improved when we used A-PoD for deconvolution.

## METHODS

### Image pre-processing

The image of the 1 μm bead was interpolated 2 times along the optical axis direction, and the retina image was interpolated 6 times in all directions. The 3D live cell images were interpolated 10 times along the optical axis direction The measured DO-SRS images were resampled before deconvolution. For all resampling process, Fourier interpolation code about f-SOFI was used.(66) In order to increase the signal-to-noise ratio, the PURE denoise filter was used 10 times to reduce noise in imaging standard bead; and automatic correction of sCMOS-related noise (ACsN) algorithm, and ACsN was used for the retina image.(67, 68)

### A-PoD algorithm

The A-PoD algorithm described in the paper was newly implemented for SRS analysis. We adopted the Adam solver (71) as the optimization method and used a gradient algorithm instead of genetic algorithm. The optimization method was changed to a gradient descent algorithm from genetic algorithm. The optimization method used is the Adam solver(69). Because the variables of A-PoD are positions of each virtual emitters, the numbers are set to the address value of the pixel. Therefore, all of these numbers have integer values, and for this, the gradient equation of the Adam solver was modified as follows.

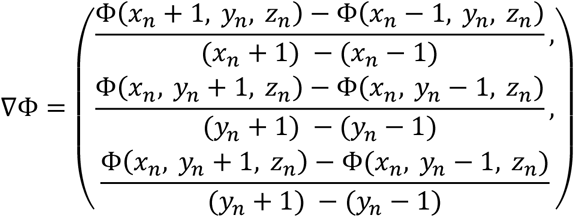

Here, Φ is an objective function for deconvolution of 3D image.

PSFs for deconvolution processes were simulated using the PSF generator in ImageJ plugin according to the physical conditions of each measurement.(20, 70) In order to efficiently process a 3D image, the image was deconvolved by dividing the image into several pieces as used in the SPIDER algorithm(23). A-PoD was implemented using Tensorflow 1.15 and Python 3.6. The number of virtual emitters used was manually controlled under the condition that the image contrast improved. All calculations were performed on a Xeon W-2145 CPU, 64 GB RAM, and NVIDIA Quadro P4000 GPU.

### Lipid droplet analysis

After deconvolution, individual lipid droplets were counted with 3D objects counter in ImageJ. Based on the information of position, volume, surface area, and mean-distance, we prepared the plots in Fig.3, 6, Extended Data Fig. 5, and 6. After detection of individual lipid droplets, we mapped the surface area to volume ratio using home-built Matlab code.

### Standard Beads

Colloid suspension of 270 nm diameter polystyrene beads with a solid content of 1.0wt.% (ThermoScientific) was used in the following experiments. To tailor the suspension for the CAPA experiments, the colloidal solution was further diluted 10-fold to a 0.1wt.% concentration (9.33×1010part/ml) using deionized water.

### Retinal section preparation

Human retina tissue sections were obtained from a donor (age 83) (San Diego Eye Bank, CA, USA) with appropriate consent from the San Diego Eye bank and following a protocol approved by the University of California, San Diego Human Research Protection Program. The donor had have no history of eye disease, diabetes, or any neurological diseases. Following fixation, the retina was process for cryostat sections (12 μm) and stored at –80 °C. Frozen sections were defrosted (10 min, RT) and washed with1X PBS 3 times, for 10 mins each time and then sandwiched between a 170nm coverslip and a glass slide with PBS solution. The coverslips were sealed with nail polish.

### MCF-7 breast cancer cell

MCF-7 cells were cultured in DMEM growth media supplemented with 10mg/L insulin (Sigma Aldrich, St. Louis, MO), 1% v/v Penicillin-Streptomycin mix (Fisher Scientific, Waltham, MA), and 5% v/v heat inactivated FBS on #1 thickness cover-glass (GG12-Laminin, Neuvitro) for 48hrs. Cells were fixed with 4%v/v PFA solution for 15 mins and then mounted on 1mm thick glass slides.

### HEK293 cells

HEK293 cells were stably transfected with a plasmid expressing monomeric red fluorescent protein containing a mitochondrial targeting sequence (Mito-Red)(71). Cells were cultured on coverglasses in 24 well cell culture dishes at 37°C (5% CO2) in DMEM supplemented with 10% fetal bovine serum (FBS; Atlanta Biological) and 1% penicillin/streptomycin (Fisher Scientific). Cells were fixed with 4% paraformaldehyde (PFA) in PBS. Following washes with PBS, the coverglasses were mounted in PBS before imaging.

### HeLa cell

HeLa cells were cultured in Dulbecco’ s modified Eagles’ medium (DMEM), supplemented with 10% fetal bovine serum (FBS) and 1% penicillin/streptomycin (Fisher Scientific, Waltham, MA), and incubated with 5% CO2 at 37°C. After passaging at 80% confluence, cells were seeded at a concentration of 2×10^5^/mL onto coverglass in a 24-well plate. DMEM with 0.5% FBS and 1% penicillin/streptomycin was used to synchronize the cells for 8 hours. The media was then changed to 50% (v/v) heavy water (D_2_O) and treatment media as described below.

For the excess aromatic amino acids condition, phenylalanine and tryptophan were increased as two separate test conditions at a 15x concentration. L-phenylalanine powder (SLCF3873, Sigma Aldrich) and L-tryptophan powder (SLCF2559, Sigma Aldrich) were added to DMEM for the excess groups. Cells were then cultured for 36 hours. Next, the cells were gently rinsed with 1x PBS with Calcium and Magnesium ions at 37°C (Fisher Scientific, 14040216), and fixed in 4% methanol-free PFA solution (VWR, 15713-S) for 15 minutes. The cover glass was finally mounted on the cleaned 1mm thick glass microscope slides with 120 μm spacers filled with 1x PBS for imaging and spectroscopy. These samples are stored at 4°C when not in use.

### Drosophila

The *w*^*111*8^parent flies were raised in vials containing the standard food (Bloomington cornmeal-yeast-sugar recipe) at 25°C in a controlled light (12/12-h light/dark cycle) and humidity (>70%) environment for several generations. The embryos from the young females (∼7 day aged) were collected in a 4 h window to synchronize larval development. Two groups of 10-15 1^st^ instar larvae were put into vials containing 20% D_2_O labeled standard food (100g yeast, 50g sucrose, 5g agar per liter) and 3x high glucose food (100g yeast, 150g sucrose, 5g agar per liter), respectively. The larvae were allowed to develop until wandering 3^rd^ instar and then brains were dissected in PBS and fixed in 4% formaldehyde for 21 min at room temperature (RT). After fixation, brains were washed four times with PBS in glass wells and were then sandwiched between a coverglass and the slide with PBS solution. To prevent the tissue drying, nail polish was used to seal the surrounding of the cover glass.

### STORM imaging

The mouse hippocampal neuronal culture and immunostaining were performed as described previously (64). The STORM imaging(72) was performed on a custom inverted microscope (Applied Scientific Imaging) with a 60x Nikon objective (MRD01605). A custom Lumencor Celesta system was used to illuminate the sample. An ∼1W 640 nm laser line was used to image a hippocampal neuron immunostained using anto-beta II spectrin antibody conjugated to the Alexa-647 dyes conjugated to the spectrin antibody and a ∼200mW 405 laser line was used to stimulate the cycling of the dyes. The Teledyne Kinetix camera was used to for imaging at 50Hz. The other imaging conditions and the parameters for the DAOSTORM fitting and processing were set as described previously.(73)

The neuron culture was performed as described previously.(64)

### SRS microscopy

A custom-built upright laser-scanning microscope (Olympus) with a 25x water objective (XLPLN, WMP2, 1.05 NA, Olympus) was applied for near-IR throughput. Synchronized pulsed pump beam (tunable 720–990 nm wavelength, 5–6 ps pulse width, and 80 MHz repetition rate) and Stokes (wavelength at 1032 nm, 6 ps pulse width, and 80MHz repetition rate) were supplied by a picoEmerald system (Applied Physics & Electronics) and coupled into the microscope. The pump and Stokes beams were collected in transmission by a high NA oil condenser (1.4 NA). A high O.D. shortpass filter (950 nm, Thorlabs) was used that would completely block the Stokes beam and transmit the pump beam only onto a Si photodiode for detecting the stimulated Raman loss signal. The output current from the photodiode was terminated, filtered, and demodulated by a lock-in amplifier at 20MHz. The demodulated signal was fed into the FV3000 software module FV-OSR (Olympus) to form image during laser scanning. All images obtained were 512 × 512 pixels, with a dwell time 80 μs and imaging speed of ∼23 s per image.

### Fluorescence microscopy

Multiphoton fluorescence microscopy is integrated with the DIY SRS microscopy together for imaging the same region of interest with different modalities (DO-SRS signals and fluorescence signals). Mitored signal was imaged with 800nm ultrafast laser scanning two photon fluorescence excitation and detected by PMT with a 610nm band pass filter in front of it.

## ACKNOWLEDGEMENTS

We thank Drs. Kun Zhang, Wei Min, and Joerg Enderlein for helpful discussion and suggestions. Thanks to Dr. Matthew Shtrahman and Samir Saidi for providing bead samples.

## AUTHOR CONTRIBUTIONS

L. S. conceived the idea and designed the project; H. J. developed and improved A-PoD algorithm and coded it. B.B. did the STORM imaging experiments. Y. L., A.F., H.K. and P.B carried out the SRS imaging experiments and collected data with the help from L.S.; H.J. analyzed the images and generated figures with the input from L.S. and B.B. D.S. prepared human retina samples; X. C. and J. W. performed experiments using HEK293 cells. Y.L. did the Drosophila work. P.B, H.K. and A.F. did the Hela cells and breast cancer cell experiments. H. J. and L.S. wrote and revised the manuscript with the input from all other authors.

